# Crystal structure of a class II UDP-glucose—hexose-1-phosphate uridylyltransferase from *Bifidobacterium longum* involved in human milk oligosaccharide metabolism

**DOI:** 10.64898/2026.09.13.751320

**Authors:** Mayo Sato, Chihaya Yamada, Mamoru Nishimoto, Motomitsu Kitaoka, Shinya Fushinobu

## Abstract

Infant-associated bifidobacteria utilize lacto-*N*-biose I (LNB) and galacto-*N*-biose (GNB), major components of human milk oligosaccharides and intestinal mucin, respectively, through the GNB/LNB metabolic pathway. A UDP-glucose–hexose-1-phosphate uridylyltransferase (GalT) from *Bifidobacterium longum* JCM 1217 (BlGalT2) catalyzes a key step in this pathway. BlGalT2 belongs to class II GalTs within the histidine triad (HIT) superfamily, a rare protein group distinct from canonical class I GalTs. Here, we determined the crystal structure of BlGalT2 at 2.25 Å resolution, providing the first three-dimensional structure of a class II GalTs. BlGalT2 is a monomeric enzyme with an architecture distinct from canonical class I GalT, comprising characteristic HIT1 and HIT2 catalytic core subdomains and an extended auxiliary domain that forms a self-contained active site. Kinetic analysis revealed a strong preference for GalNAc-1P over Gal-1P, with a 50-fold lower *K*_m_ for GalNAc-1P, consistent with a hydrophobic pocket accommodating the *N*-acetyl group. Metal analysis and activity measurements indicated that Zn^2+^ is required for full activity, and structural prediction suggested its coordination by residues including the first histidine of the HIT motif. AlphaFold3 prediction and mutational analysis identified residues involved in nucleotide-sugar recognition and supported a Ping-Pong reaction mechanism involving a covalent enzyme–uridine monophosphate intermediate. These findings reveal how class II GalTs have evolved a unique structural framework and substrate specificity for *N*-acetylated sugars. Moreover, the structure of BlGalT2 fills the final missing gap in bifidobacterial HMO metabolism and provides molecular insight into the adaptation of bifidobacteria to host-derived glycans.

## INTRODUCTION

Bifidobacteria confer various beneficial effects on human health as symbiotic gut bacteria and are widely used as probiotics [1]. In the early life stages of breast-fed infants, bifidobacterium species become major constituents of the gut microbiota [2]. This bifidus-flora formation can be attributed to the presence of the specific degradation pathway for human milk oligosaccharides (HMOs) in the infant-associated bifidobacteria [3,4]. Lacto-*N*-biose I (Galβ1,3GlcNAc; LNB) is a major building block of HMOs, and infant-associated bifidobacteria, namely *Bifidobacterium longum* subsp. *longum* (*B. longum*), *Bifidobacterium longum* subsp. *infantis*, *Bifidobacterium bifidum*, and *Bifidobacterium breve*, possess a specific degradation pathway for this disaccharide. This pathway can also effectively degrade galacto-*N*-biose (Galβ1,3GalNAc; GNB), which comprises one of the core structures of intestinal mucin glycoproteins. Therefore, this pathway is called GNB/LNB pathway [5]. Enzymes involved in the GNB/LNB pathway of *B. longum* JCM 1217 have been fully characterized: a GNB/LNB-specific ATP-binding cassette (ABC)-type oligosaccharide transporter (EC 7.5.2.2; locus tag = BLLJ_1624-26) [6], GNB/LNB phosphorylase (EC 2.4.1.211; BLLJ_1623) [7], *N*-acetylhexosamine 1-kinase (EC 2.7.1.162; BLLJ_1622), UDP-glucose—hexose-1-phosphate uridylyltransferase (BlGalT2; BLLJ_1621), and UDP-glucose /UDP-*N*-acetylglucosamine 4-epimerase (EC 5.1.3.2/EC 5.1.3.7; BLLJ_1620) [8]. The GNB/LNB pathway is an energy-saving variation of the well-known Leloir pathway, which is responsible for galactose metabolism in most organisms, because the phosphorylase attaches a phosphate to Gal in LNB or GNB without consuming ATP [9]. BlGalT2 catalyzes the following reactions to convert *galacto*-configured compounds derived from LNB and GNB into *gluco*-configured compounds that are metabolized by glycolysis and amino sugar metabolism.

Glc-1P + UDP-Gal ⇔ Gal-1P + UDP-Glc (EC 2.7.7.12)

GlcNAc-1P + UDP-GalNAc ⇔ GalNAc-1P + UDP-GlcNAc (EC 2.7.7.-)

In the GNB/LNB pathway, the crystal structures of the solute-binding protein of the ABC-type transporter (GL-BP) [6], GNB/LNB phosphorylase (GLNBP) [10], *N*-acetylhexosamine 1-kinase (NahK) [11], and UDP-glucose 4-epimerase (bGalE) [12] have been reported. However, only the three-dimensional structure of BlGalT2 remains unknown.

UDP-glucose—hexose-1-phosphate uridylyltransferases (GalTs), including BlGalT2, belong to the histidine triad (HIT) protein superfamily that features a conserved HφHφHφφ motif (φ = hydrophobic amino acid) [13]. The HIT superfamily enzymes utilize Ping-Pong mechanism because the second His residue in the HIT motif nucleophilically attacks the α-phosphate of UDP-sugar to form a covalent enzyme-uridine monophosphate (UMP) intermediate (E-UMP), and UMP is subsequently transferred to an acceptor substrate. The HIT superfamily is divided into four families: Hint, Fhit, Aprataxin and GalT [13]. GalT family proteins have a modified HIT motif, HXHXQφφ and are further classified into three groups: GalT, Ap_4_A phosphorylase, and adenylylsulfate:phosphate adenylyltransferase groups. In the InterPro classification (https://www.ebi.ac.uk/interpro) [14], members of the GalT group are further divided into classes I and II, which share very low amino acid sequence identity with each other (<15%). Class I GalTs (IPR001937) are widely present in bacteria, eukaryotes, and archaea, whereas class II GalTs (IPR000766) are very rare protein family that is found only in the bacterial phyla Bacillota (formerly Firmicutes) and Actinomycetota (formerly Actinobacteria) (Fig. 1). There are a few reports on class II GalTs, and only enzymes from *Lactobacillus helveticus* [15], *B. longum* [8], and *B. bifidum* [16] have been biochemically characterized. Infant-associated bifidobacteria have both class I and II GalT genes (*galT1* and *galT2*) in the genomes, and the latter is involved in the GNB/LNB pathway [16]. Sequence identity between the two GalTs from *B. bifidum* ATCC 29521 (BbGalT1 and BbGalT2) was only 12.1% and their specific activities toward Gal-1P and UDP-Glc are 93.1 and 0.261 U/mg, respectively. The most remarkable difference between BbGalT1 and BbGalT2 is that the former can only use Gal-1P/Glc-1P and UDP-Glc/UDP-Gal as substrates, while the latter can also use GalNAc-1P/GlcNAc-1P and UDP-GlcNAc/UDP-GalNAc.

**Figure 1.**
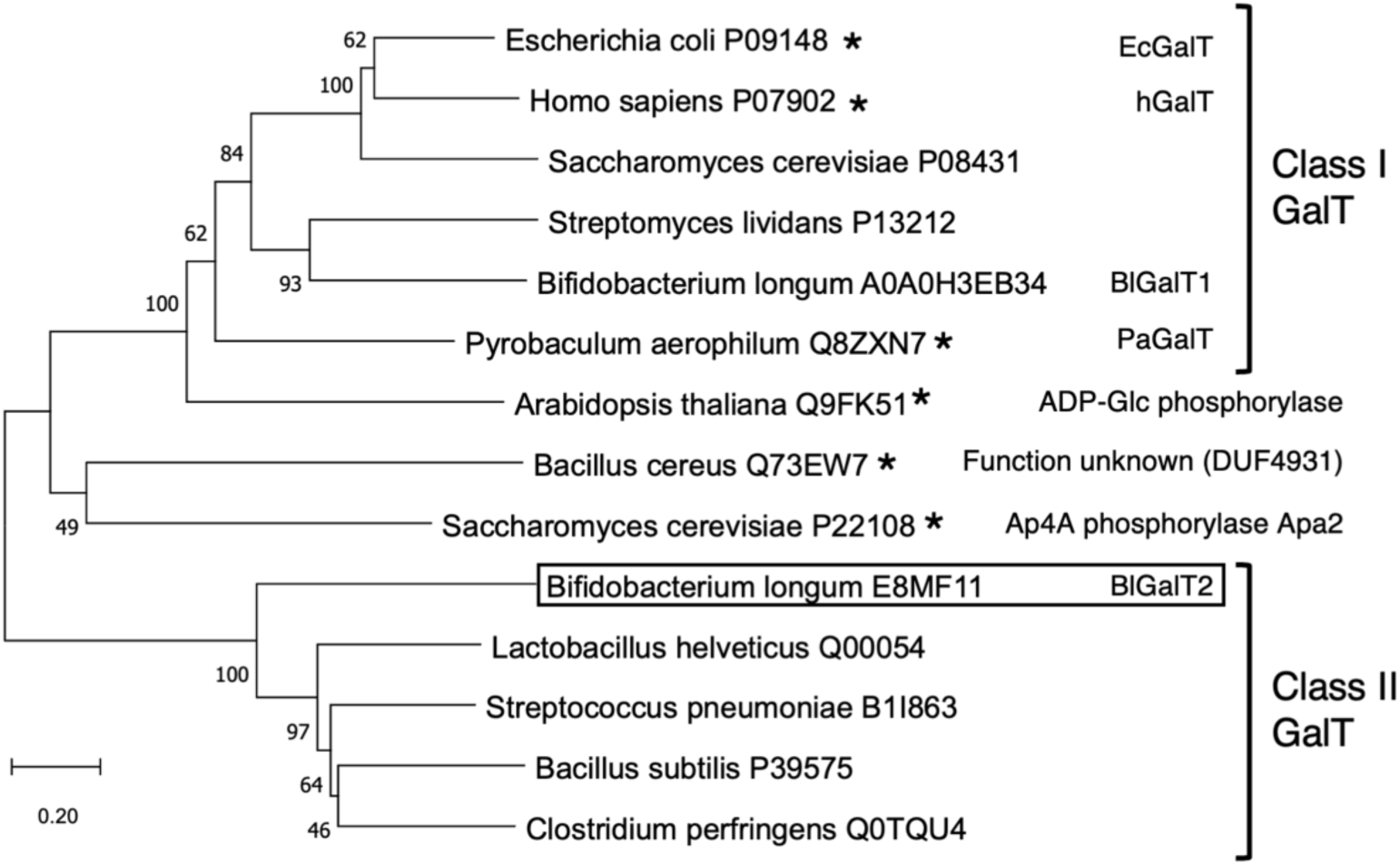
A phylogenetic tree of GalTs and structurally related enzymes. The UniProt entry identifier is shown after the source organism name. Asterisks (*) indicate structurally determined enzymes. The optimal tree was generated using the Neighbor-Joining method. Bootstrap values of 100 replications are indicated at the branches. The scale indicates branch lengths measures in the number of substitutions per site.

Here we report the crystal structure of BlGalT2 from *B. longum* JCM 1217 (BLLJ_1621/LnpC) as the first three-dimensional structure of class II GalT. We also revealed that BlGalT2 prefers GalNAc-1P over Gal-1P through kinetic analysis. This report fills the final missing structural piece in understanding HMO degradation by infant gut-associated bifidobacteria.

## MATERIALS AND METHODS

### Protein production and purification

A pET19b(+)-based vector encoding N-terminally His_10_-tagged BlGalT2 (pET19b(+)-*galT_Bl_*, residues 1–515) was constructed from the pET30-based expression vector carrying the *lnpC* gene of *B. longum* JCM 1217 [8]. The primers listed in Table S1 were used for vector to insert a TEV protease site after the His_10_-tag. The plasmid was introduced into *Escherichia coli* BL21-CodonPlus (DE3)-RIL (Stratagene, La Jolla, CA) for protein expression. The transformants were precultured at 37°C for 4 h in lysogeny broth medium containing 100 mg/L ampicillin and 30 mg/L chloramphenicol. A 5 mL portion of the culture was inoculated into 200 mL of fresh lysogeny broth medium until the optical density at 600 nm reached 0.6. Protein production was induced with 0.1 mM isopropyl 1-thio-β-D-galactopyranoside at 25°C for 10 h. The cells were harvested by centrifugation and suspended in 50 mM Tris-HCl (pH 7.8). Cell extracts were obtained by sonication followed by centrifugation to remove cell debris. The supernatant was applied to a HisTrap FF crude column (Cytiva, Marlborough, MA, USA) pre-equilibrated with 50 mM Tris-HCl (pH 7.8) containing 100 mM NaCl, and the column was washed with 30 mM imidazole and then eluted with 400 mM imidazole at a flow rate of 4 mL/min at 4°C. The purified protein obtained at this step was used for activity assays. Because removal of the His_10_-tag was required for crystallization, further purification was performed through the following steps. Imidazole was removed from the HisTrap elution peak fractions and the buffer was exchanged into 20 mM Tris-HCl (pH 7.8) using an Amicon Ultra centrifugal filter device (30 kDa MWCO; Merck, Darmstadt, Germany). The protein concentration was adjusted to 1 mg/mL with the same buffer. A TEV protease solution was added at a 1:50 (v/v) ratio, and the reaction mixture was incubated overnight at 4 °C. The mixture was then loaded onto a HisTrap column as described in the previous step, and the flow-through fraction was collected and concentrated using an Amicon Ultra centrifugal filter device (30 kDa MWCO, Merck). The protein was then subjected to gel filtration on a HiLoad 16/600 Superdex 200 pg column (Cytiva) pre-equilibrated with 20 mM Tris-HCl (pH 7.8) containing 150 mM NaCl at a flow rate of 1 mL/min at 4°C. The purified protein was concentrated and buffer-exchanged to 10 mM Tris-HCl (pH 7.8) using an Amicon Ultra centrifugal filter device (30 kDa MWCO, Merck). Protein concentrations were determined by measuring the absorbance at 280 nm on a NanoDrop Lite Plus spectrophotometer (Thermo Fisher Scientific). The molar extinction coefficient was calculated from the amino acid sequence using the ProtParam tool on ExPASy (https://web.expasy.org/protparam/). To produce a selenomethionine (SeMet) derivative of BlGalT2, *E. coli* cells harboring the expression plasmid were cultivated in a modified M9 medium containing SeMet and inhibitors of methionine biosynthesis [17]. The SeMet derivative of BlGalT2 was purified using the same method as the native protein.

### Metal content analysis

Inductively coupled plasma mass spectrometry (ICP-MS) measurements were performed using an ICPS-8100 (Shimadzu, Kyoto, Japan) spectrometer. All solutions for ICP-MS analysis were prepared with metal-free ultrapure water. For calibration curve preparation, 1.5 g of 0.08 N HNO_3_ solution containing 2 ppb In (indium) solution was accurately weighed. Stock solutions of each metal (1000 ppm) were weighed out (1.5 mg each) to prepare individual 1 ppm (1000 ppb) solutions. These solutions were further diluted with the 0.08 N HNO_3_ + 2 ppb In solution to final concentrations of 100 ppb for Ca^2+^ and Mg^2+^ and 20 ppb for the other metals. Serial 1:5 dilutions were prepared to yield five calibration standards. To eliminate carryover errors from the buffer, equal amounts of sample were used regardless of sample concentration. Specifically, 50 µL of 2% HNO_3_ was placed at the bottom of a 15 mL tube, mixed with 30 µL of sample solution, and thoroughly vortexed. To this mixture, 5 mL of the 0.08 N HNO_3_ + 2 ppb In solution was added, and the mixture was filtered through a 0.22 µm membrane. Each sample was serially diluted 1:5 to perform measurements across three dilution series per sample. Blank samples containing buffer only were prepared in the same manner for each dilution series and subtracted from the sample readings, which also allowed verification of dilution-induced errors. Finally, the molar ratio of metal to protein was determined from the measurement results, accounting for dilution factors, the protein molecular weight, and the atomic weights of the metals. Protein concentrations were determined using a BCA protein assay kit (Thermo Fisher Scientific, Waltham, MA, USA) with bovine serum albumin as a standard.

### Enzyme assay and kinetic analysis

The enzymatic activity of BlGalT was assayed using UDP-Glc as the nucleotide sugar substrate and Gal-1P or GalNAc-1P as the phosphorylated sugar substrate. The concentration of NADPH generated by coupling the reaction-produced Glc-1P to a phosphoglucomutase and glucose-6-phosphate dehydrogenase reaction was measured by monitoring the increase in absorbance at 340 nm. The standard assay mixture (50 µL) contained 50 mM HEPES-NaOH buffer (pH 8.0), 2 µM glucose 1,6-bisphosphate (Sigma-Aldrich), 0.25 mM NADP^+^ (Oriental Yeast, Tokyo, Japan), 0.25 U of phosphoglucomutase from rabbit muscle (Sigma-Aldrich), 0.25 U of glucose-6-phosphate dehydrogenase from yeast (Oriental Yeast), 5 mM MgCl_2_, 1 mM UDP-Glc (Toyobo, Osaka, Japan), 1 mM Gal-1P or GalNAc-1P (Biosynth, Staad, Switzerland), and an appropriate amount of the enzyme. The reaction was started by mixing the BlGalT2 protein solution (10 μL) and the remaining assay mixture (40 μL), both of which had been preincubated at 35°C for 5 min. The reaction was monitored using a Benchmark Plus microplate reader (Bio-Rad, Hercules, CA, USA) with a 96-well flat-bottomed transparent microplate #9018 (Corning, Corning, NY, USA) at 35°C. The absorbance increase at 340 nm was monitored for up to 20 min at 20 s intervals, and the enzyme concentration was set to observe a linear absorbance increase (initial rate) within this time frame. The following substrate concentrations were used for the determination of kinetic constants: 0.125 to 4.0 mM UDP-Glc, 0.50 to 5.0 mM Gal-1P, and 0.125 to 1.0 mM GalNAc-1P. A standard curve was created using 0 to 0.50 mM NADPH. Kinetic analysis was performed using the Enzyme-Kinetic Calculator [18].

To measure metal concentration dependence, standard assay mixtures supplemented with ZnCl_2_ to a final concentration of 0.001–0.1 mM were pre-incubated at 35 °C for 10 min, after which the enzyme solution was added to initiate activity measurements. To prepare the metal-depleted protein, the enzyme was incubated in 10 mM Tris-HCl (pH 7.8) containing 10 mM Na-ethylenediaminetetraacetic acid (EDTA) at 4 °C for 30 min. Excess EDTA was subsequently removed by exchanging the buffer to 10 mM Tris-HCl (pH 7.8) using an Amicon Ultra centrifugal filter device (30 kDa MWCO, Merck).

### Crystallography and bioinformatic analyses

Crystals of native BlGalT2 and its SeMet derivative were obtained at 4°C using the sitting drop vapor diffusion method. Specifically, 0.5 µL protein solution (10 mg/mL) was mixed with an equal volume of a reservoir solution containing 0.5 M LiCl, 0.1 M Na-citrate (pH 5.0), and 15% PEG 3350. Crystals were cryoprotected in the reservoir solution supplemented with 20% (w/v) ethylene glycol and flash-cooled at 100 K in a stream of nitrogen gas. X-ray diffraction data were collected at the beamlines of the Photon Factory of the High Energy Accelerator Research Organization (KEK, Tsukuba, Japan) and SPring-8 (Hyogo, Japan). Data processing was performed using XDS [19] and Aimless [20]. The crystal structure of BlGalT2 was solved by single-wavelength anomalous dispersion using SeMet-derivative. Initial phase calculation and automated model building were performed using Phenix [21]. Manual model rebuilding and refinement were carried out using Coot [22] and Refmac5 [23], respectively. Atomic coordinates and structure factors for the crystal structure have been deposited in the Protein Data Bank Japan (https://pdbj.org) under accession code 45PI.

Phylogenetic analysis was performed using MEGA 11.0.13 [24]. The protein sequences were aligned using MUSCLE [25]. Protein structure prediction was performed using AlphaFold3 [26] installed on a PC with the source code (https://github.com/google-deepmind/alphafold3) and the model parameters distributed by Google DeepMind. Molecular graphics images were prepared using PyMOL Schrödinger LLC, New York, NY, USA).

### Construction of mutant enzymes

BlGalT2 mutant enzymes were constructed using the QuikChange Site-Directed Mutagenesis Kit (Agilent Technologies) with the pET19b(+)-*galT_Bl_* plasmid as a template. The primers listed in Table S1 and their complementary strands were used. The entire ORF sequence was verified to ensure that no base changes other than the designed mutations had occurred. The mutant enzymes were expressed, purified, and assayed by procedures similar to those described above.

## RESULTS

### Metal content determination, metal dependence, and kinetic analysis

The recombinant BlGalT2 protein was heterologously expressed in *E. coli* and purified (Fig. S1). The molecular mass of BlGalT2 deduced from its amino acid sequence, estimated by SDS-PAGE, and determined by calibrated gel filtration chromatography was 56.7, 57, and 58 kDa, respectively, indicating that it is monomeric in solution.

Structurally characterized class I GalTs possess an essential structural zinc-binding site conserved near the active site [27–29]. GalTs from *E. coli* (EcGalT) and *Pyrobaculum aerophilum* (PaGalT) additionally contain a nonessential structural iron-binding site, which is not conserved in the human enzyme. Therefore, we analyzed the metal content of BlGalT2 (Table S2). The purified recombinant BlGalT2 protein contained 1.06 ± 0.13 mol/mol Zn^2+^, 0.30 ± 0.11 mol/mol Fe^2+^, and 0.04 ± 0.00 mol/mol Ni^2+^, with no detectable amounts of other major metal ions (Mg^2+^, Ca^2+^, Mn^2+^, Co^2+^, Cu^2+^, V^2+^, and Cr^2+^). The enzyme activity was unaffected by the addition of Zn^2+^ at low concentrations but was inhibited at 0.1 mM Zn^2+^ (Table 1). Treatment with EDTA reduced the specific activity to approximately 75% of the untreated level, and the activity was restored by the addition of 0.1 mM Zn^2+^. These results suggest that BlGalT2 contains a catalytically essential zinc-binding site.

**Table 1.** Effect of Zn^2+^ addition and EDTA treatment on the activity of BlGalT2.

| ZnCl (mM) | Specific activity (U/mg) | Specific activity after EDTA treatment (U/mg) |
| --- | --- | --- |
| 0 | $0.61 \pm 0.10$ | $0.46 \pm 0.1$ |
| 0.001 | 0.60 | 0.29 |
| 0.01 | 0.60 | 0.34 |
| 0.1 | 0.25 | 0.63 |

The activity toward 1 mM UDP-Glc and 1 mM Gal-1P was measured at 35°C in 50 mM HEPES-NaOH buffer (pH 8.0).

We previously reported the specific activities of BlGalT2 toward Gal-1P and GalNAc-1P under defined conditions [8], but kinetic analyses were not performed. Here, we conducted steady-state kinetic analyses of BlGalT2 using UDP-Glc as the donor substrate and Gal-1P or GalNAc-1P as the acceptor substrates. When the initial reaction rates were plotted against the concentration of one substrate while the concentration of the other substrate was fixed, typical Ping-Pong Bi-Bi mechanism patterns were observed in Lineweaver-Burk-type double-reciprocal plots (Fig. 2). The kinetic parameters for each substrate were determined by nonlinear global fitting of these data (Fig. S2 and Table 2). For the two acceptor substrates (Gal-1P and GalNAc-1P), the *K*_m_ values for UDP-Glc and the *k*_cat_ values were comparable. In contrast, the *K*_m_ value for GalNAc-1P was approximately 50-fold lower than that for Gal-1P, indicating that BlGalT2 has a substantially higher affinity for substrates containing an *N*-acetyl group. These results indicate that BlGalT2 is more appropriately designated as UDP-GlcNAc–*N*-acetylhexosamine-1-phosphate uridylyltransferase.

**Figure 2.**
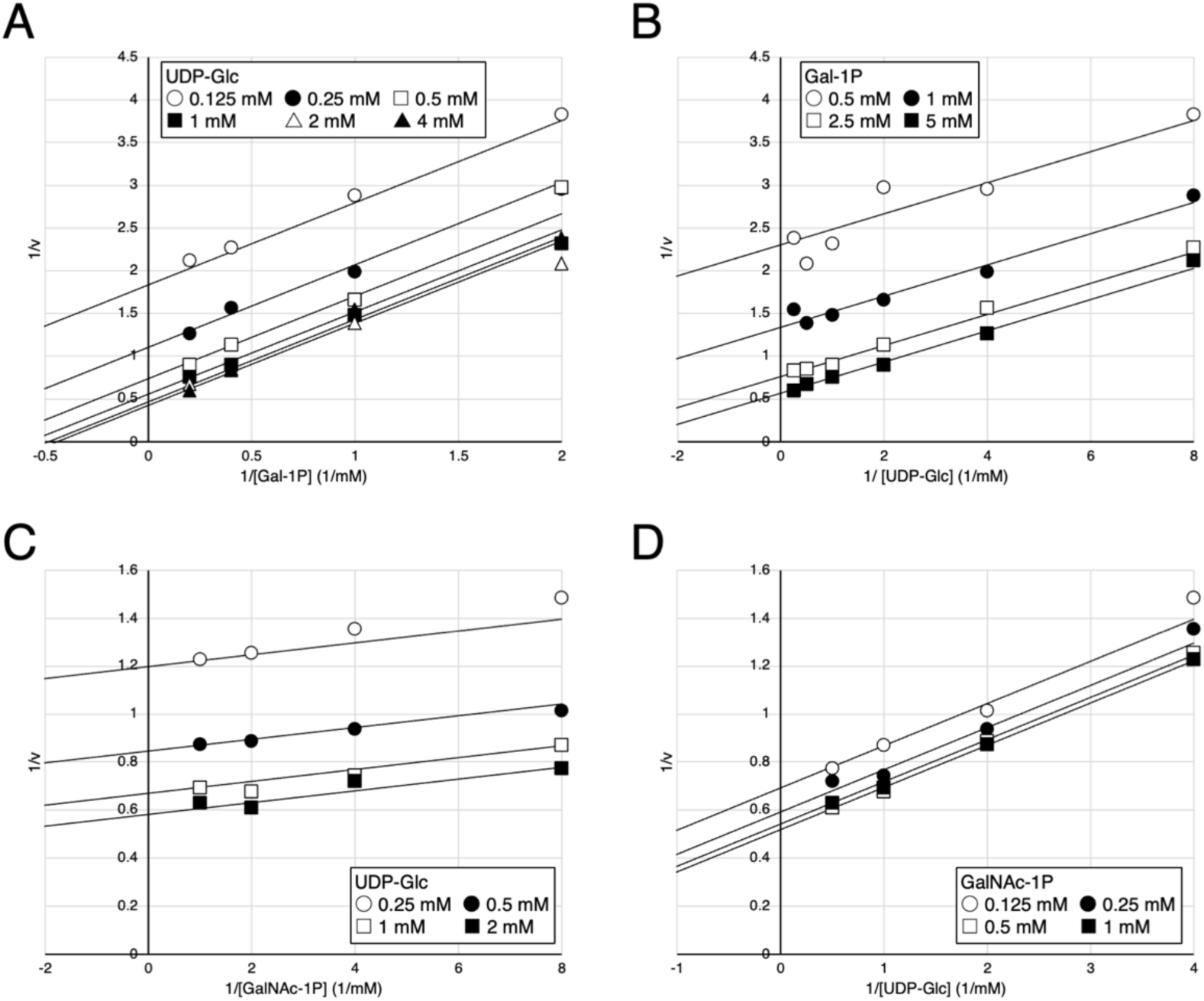
Double-reciprocal plots showing the Ping-Pong Bi-Bi kinetics of BlGalT2. (A and B) Initial velocity plots for the reactions with UDP-Glc and Gal-1P. (C and D) Initial velocity plots for the reactions with UDP-Glc and GalNAc-1P. Lines represent the global nonlinear curve fitting of the substrate-velocity (*v* vs [*S*]) data shown Figure S2.

**Table 2.**
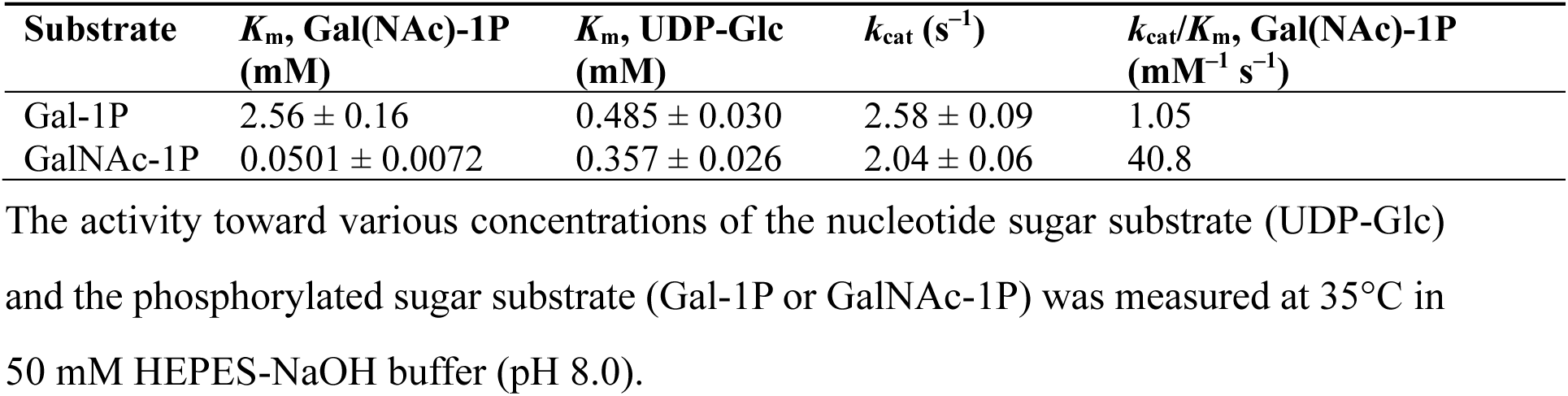
Kinetic parameters of BlGalT2.

### Crystal structure and comparison with structural homologs

BlGalT2 was crystallized, and its ligand-free form structure was determined at 2.25 Å resolution (Table S3). Despite soaking and co-crystallization attempts with UDP-sugar substrates, phosphorylated sugar substrates, and various monosaccharides, no complex structures were obtained. The asymmetric unit of the ligand-free crystal contained two BlGalT2 molecules, designated chains A and B (Fig. S3). For chain A, residues 4–48, 75– 172, 209–281, and 285–515 were modeled (Fig. 3A), whereas for chain B, residues 4–48, 75–172, 209–281, and 286–515 were modeled. Each chain contained one chloride ion (Cl^−^). Because this chloride-binding site is not conserved in structural homologs and is located far from the active site, its functional significance remains unclear. Crystal packing interface analysis using the PISA server indicated that the biological assembly of BlGalT2 is a monomer [30]. The structures of the two monomers were virtually identical, with a root-mean-square deviations (RMSD) of 0.180 Å for 410 Cα atoms. Because chain A exhibited a lower mean B-factor (Table S3), the following structural description focuses on chain A. In the catalytic HIT motif (H286, H288, and Q290), H286 showed relatively unclear electron density, whereas H288 and Q290 were well defined (Fig. 3B).

**Figure 3.**
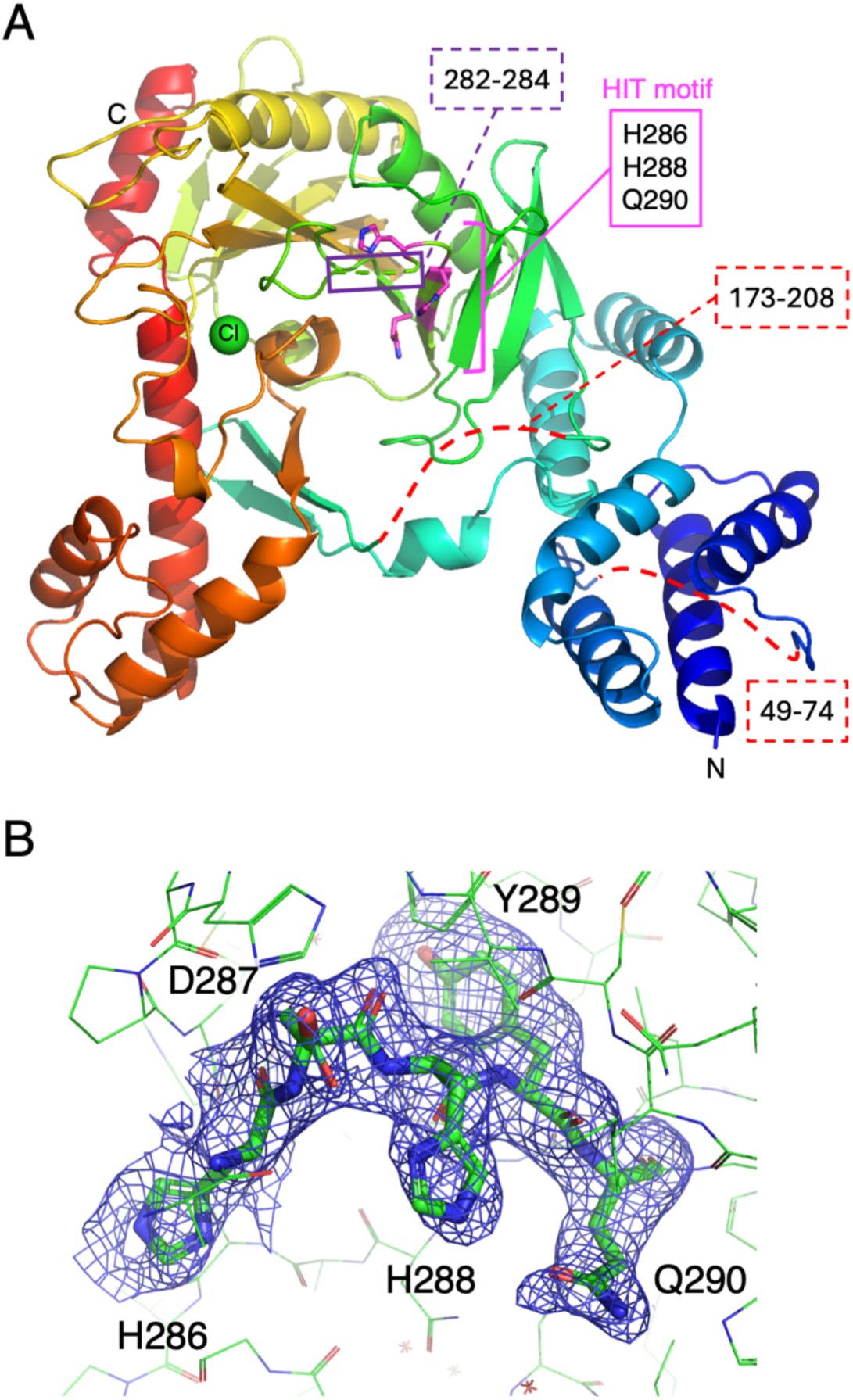
The crystal structure of BlGalT2. (A) Overall structure shown by a ribbon model in rainbow colors (blue to red from the N- to C-terminus). Side chains of the HIT motif residues are shown as magenta sticks. Disordered regions are shown by dotted lines. A Cl atom is shown as a green sphere. (B) Electron density map of the HIT motif residues. The Polder m*F*_o_-D*F*_c_ map (contoured at 1.5σ) is shown as a blue mesh.

A Dali server search [31] against three-dimensional structures deposited in the PDB (Table 3) showed that, consistent with the phylogenetic analysis based on multiple amino-acid sequence alignments (Fig. 1), BlGalT2 was more structurally similar to enzymes with different activities (Ap_4_A and ADP-Glc phosphorylases) than to class I GalTs. The highest-scoring hit was *Saccharomyces cerevisiae* Ap_4_A phosphorylase (Apa2) [32]. However, the structural similarity was not high, with a Z-score of 17.2 and an RMSD of 3.4 Å over 248 Cα atoms, and the amino acid sequence identity was less than 10% (Table 3). Apa2 consists of two domains, both of which adopt an α/β fold, namely a core catalytic HIT domain and an auxiliary domain (Fig. 4B) [32]. The core HIT domain is further divided into two subdomains, HIT1 and HIT2, and the HIT motif (HXHXQ) is located within the HIT1 subdomain. Similarly, the core catalytic domain of BlGalT2 consists of the HIT1 (residues 131–148, 209-266, and 285-295) and HIT2 (residues 267–281 and 295-395) subdomains (Fig. 4A, top). However, the auxiliary domain of BlGalT2 (residues 149–172 and 396–515) is larger than that of Apa2, and an additional helical domain (residues 4–130) is present at the N-terminus. The domain organization in the primary structure is somewhat complex, with short regions of HIT1, the auxiliary domain, and HIT2 occurring discontinuously in the central part (Fig. 4A, bottom). In the crystal structure of BlGalT2, three regions were disordered (Fig. 3): a long central region of the helical domain (residues 49–74), a long region connecting the auxiliary domain and HIT1 (residues 173–208), and a short region connecting HIT2 and HIT1 (residues 282–284). The third short, disordered region is located immediately precedes the HIT motif (residues 286–290).

**Figure 4.**
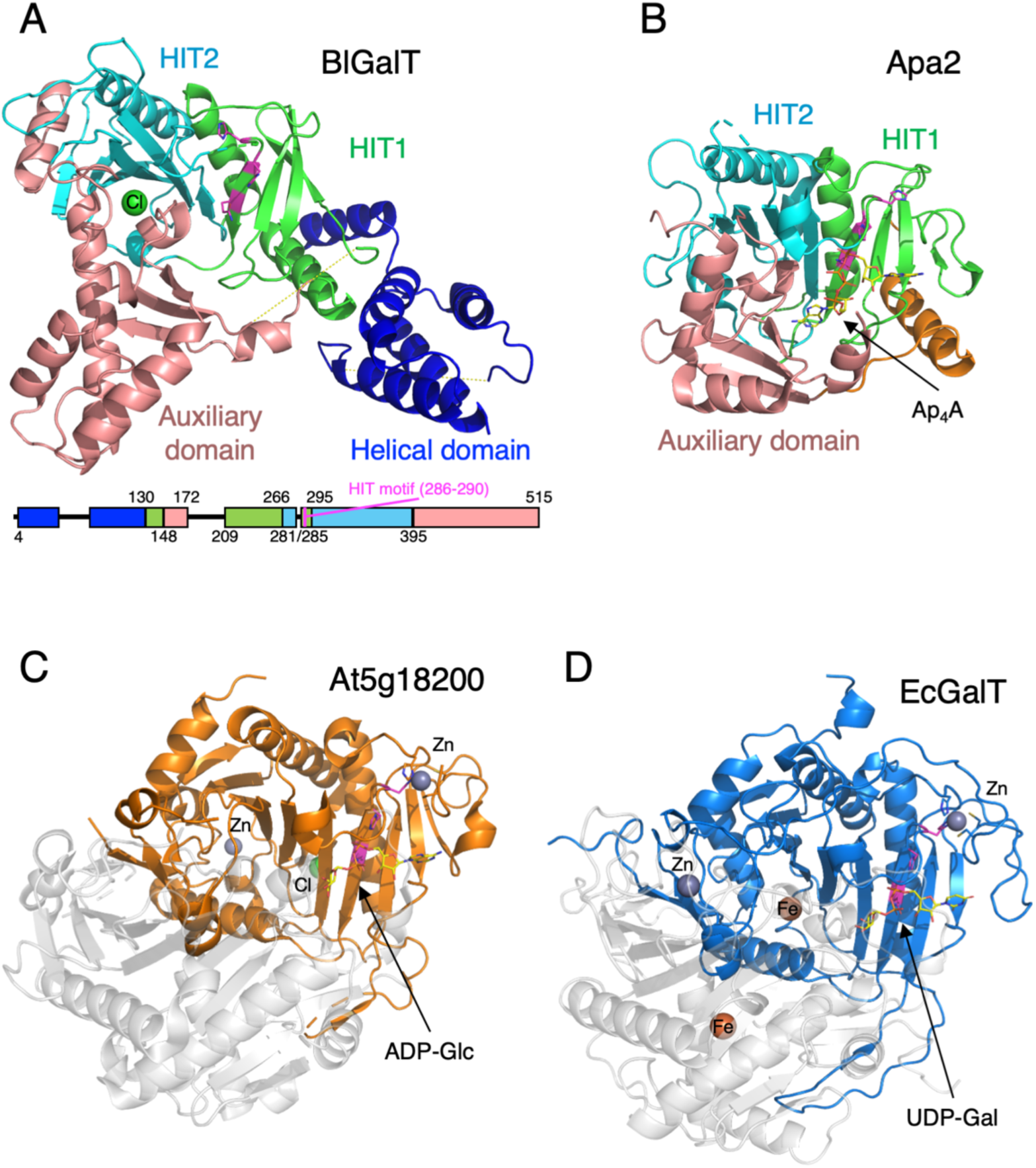
Comparison with structural homologs. (A) BlGalT2 colored by domain and subdomains: HIT1 in green, HIT2 in cyan, auxiliary domain in pink, and helical domain in blue. Side chains of the HIT motif residues are shown as magenta sticks. The domain organization in the primary structure is shown at bottom. (B) *S. cerevisiae* Ap_4_A phosphorylase Apa2. The structure of Q163H mutant complexed with Ap_4_A (PDB ID: 4I5V) is shown with the side chain of Q163 of the wild type Ap_4_A structure (PDB ID: 4I5W). (C) *A. thaliana* ADP phosphorylase At5g18200. The structure of Q186H mutant complexed with ADP-Glc (PDB ID: 2H39) is shown with the side chain of Q186 of the wild type At5g18200 (PDB ID: 1Z84). (D) *E. coli* GalT. The structure of Q166H mutant complexed with UDP-Gal (PDB ID: 1GUP) is shown with the side chain of Q166 of the wild type *E. coli* GalT (PDB ID: 1HXP). Cl, Zn, and Fe atoms are shown as green, gray, and orange spheres, respectively.

**Table 3.** Result of DALI structural similarity search.

| Protein | Organism | Activity | PDB ID (chain) | Z score | RMSD (Å) | LALI <sup>a</sup> | %ID <sup>b</sup> |
| --- | --- | --- | --- | --- | --- | --- | --- |
| Apa2 | *Saccharomyces cerevisiae* | Ap<sub>4</sub>A phosphorylase | 4I5T (A) | 17.2 | 3.4 | 248 | 9 |
| BCE0241 | *Bacillus cereus* | Unknown (DUF4931) | 4QVU (A) | 10.9 | 4.0 | 174 | 13 |
| At5g18200 | *Arabidopsis thaliana* | ADP-glucose<br>phosphorylase | 2Q4L (B) | 8.1 | 3.9 | 174 | 6 |
| PaGalT | *Pyrobaculum aerophilum* | galactose 1-phosphate<br>uridylyltransferase | 6K9Z (B) | 7.9 | 4.4 | 179 | 11 |
| hGalT | *Homo sapiens* | galactose 1-phosphate<br>uridylyltransferase | 5IN3 (B) | 7.1 | 4.5 | 173 | 8 |
| CDH | *Escherichia coli*<br>O157:H7 | CDP-diacylglycerol<br>pyrophosphatase | 2POF (B) | 7.1 | 3.5 | 150 | 8 |
| EcGalT | *Escherichia coli* | galactose 1-phosphate<br>uridylyltransferase | 1HXQ (B) | 6.9 | 4.7 | 173 | 5 |

Analyzed using the DALI server (http://ekhidna2.biocenter.helsinki.fi/dali/) [31].
<sup>a</sup> Number of aligned residues. <sup>b</sup> Sequence identity.

The second hit in the Dali structural similarity search was a functionally uncharacterized DUF4931 protein (BCE0241) from *Bacillus cereus*, and the third was *Arabidopsis thaliana* ADP-glucose phosphorylase At5g18200 (Table 3) [33]. These were followed by class I GalTs, including PaGalT [29], human GalT (hGalT) [28], and EcGalT [34]. The Z-scores of the proteins ranked third and below were below 9, indicating that their structural similarity to BlGalT2 is very low. At5g18200 and class I GalTs form homodimeric structures, and their active sites are located between the two subunits (Fig. 4C, D). Although the HIT motif characteristic of the HIT superfamily and the surrounding fold are conserved, these proteins lack an auxiliary domain. Thus, the structural analysis also demonstrates that the class II enzyme BlGalT2 differs substantially from class I GalTs.

### Active site structure predicted by AlphaFold and comparison with Ap_4_A phosphorylase and class I GalTs

Because BlGalT2 contains several disordered regions and its crystal structure was obtained in the ligand-free state, we used AlphaFold3 to predict a structure containing UDP-GalNAc and Zn^2+^ (Fig. 5A). In the predicted structure, all three disordered regions were modeled with relatively low confidence (Fig. S4). The long loop comprising residues 173–208 features a central helix that covers the active site. The short region comprising residues 282–284 was predicted to interact directly with the substrate (Fig. 5B). Zn^2+^ was predicted to be coordinated by C195, C198, H245, and H286, which is one of the histidine residues in the HIT motif.

**Figure 5.**
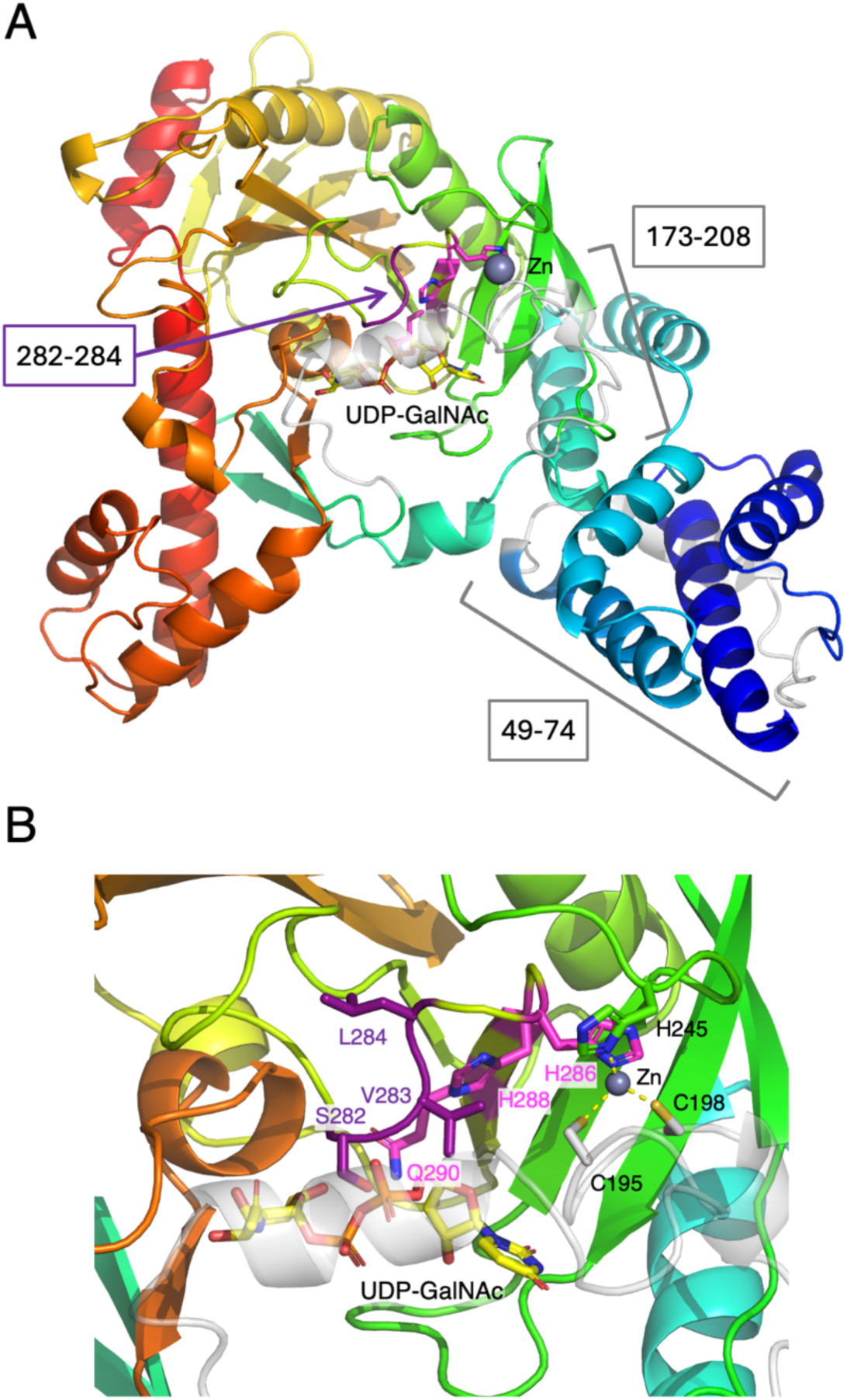
Alphafold-predicted structure of BlGalT2. (A) Overall and (B) active site structures of BlGalT2 (rainbow colors) complexed with UDP-GalNAc (yellow) and Zn^2+^ (grey). Disordered regions in the crystal structure, corresponding to residues 49–74 and 173-208, are shown as transparent gray, whereas residues 282-284 are colored purple. Side chains of the HIT motif residues are shown as magenta sticks. In (B), side chains of Zn^2+^-coordinating residues and residues 282–284 are shown as sticks.

The predicted structure allowed us to infer the substrate recognition mechanism of BlGalT2. UDP-GalNAc was predicted to form numerous polar interactions in the active site (Fig. 6A, left). Among the residues in the HIT1 subdomain, K209, Q227, and S229 recognized the adenosine moiety, whereas Y233 recognized the β-phosphate. N223 in the HIT2 subdomain did not interact with the substrate in the predicted structure of BlGalT2, despite its corresponding residue being conserved in Apa2 (N143) and EcGalT (N153) (Fig. 6B, C). Most residues comprising the active site adopted nearly identical conformations in both the crystal and predicted structures of BlGalT2. However, K209 and H286, which is one of the HIT motif residues, showed substantial deviations. This is likely because these residues are located near the termini adjacent to disordered regions in the crystal structure. In the predicted structure, the side chain of H288, which is the second histidine in the HIT motif and the putative nucleophile that attacks the substrate [35], formed a hydrogen bond with the main chain carbonyl group of H286. The side chain of H286 was predicted to coordinate Zn^2+^. In addition, four residues located in the disordered regions (K173, T196, S282, and V283) were predicted to recognize the substrate through hydrogen bonds. The pyrophosphate moiety of UDP-GalNAc was recognized by three of these residues (K173, S282, and V283), indicating that highly flexible regions play an important role in catalysis.

**Figure 6.**
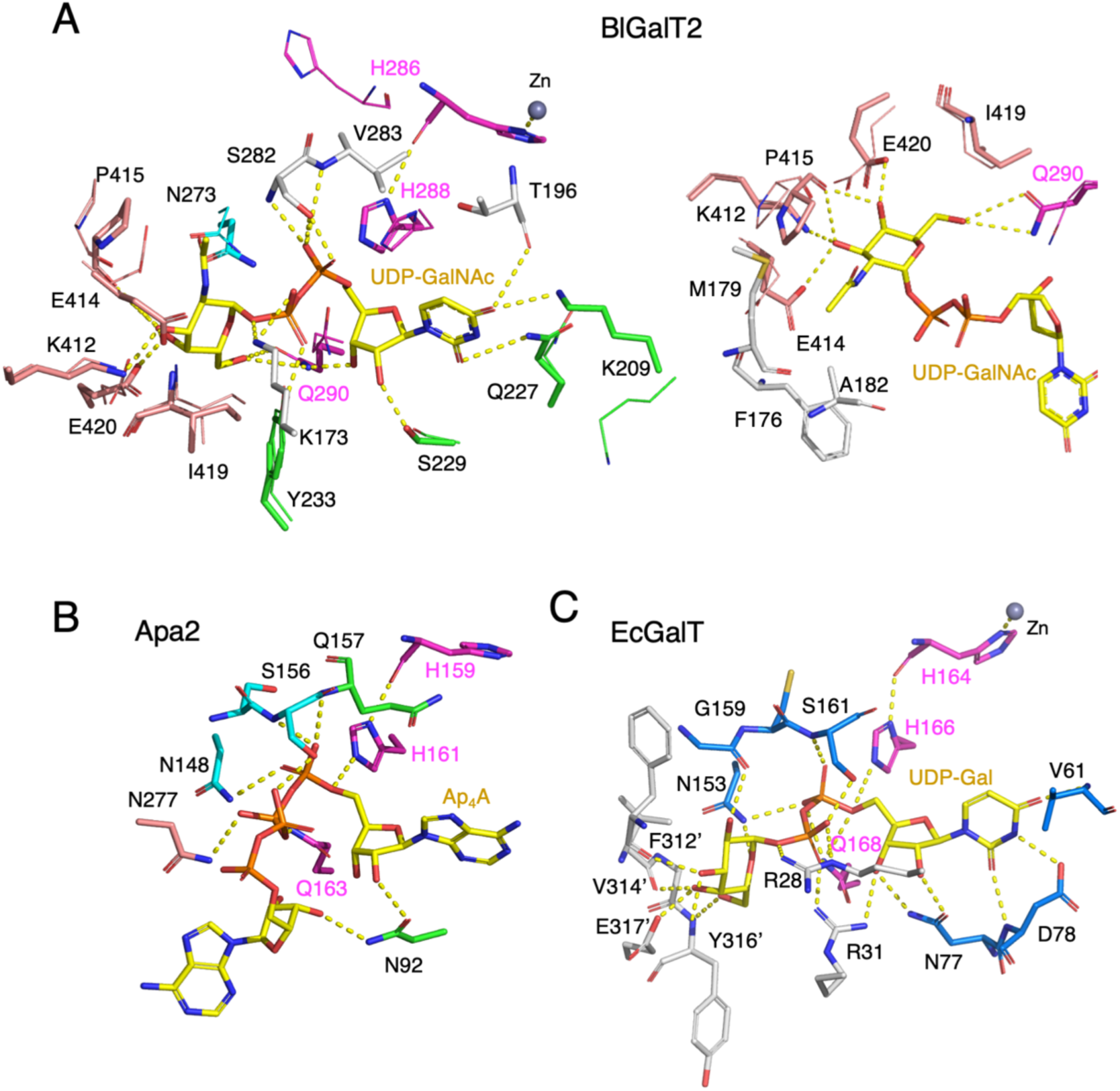
Comparison of the active sites. (A) Superimposition of the predicted (sticks) and crystal (lines) structures of BlGalT2. Residues recognizing UDP-GalNAc (left) and the GalNAc moiety (right) are shown. Residues in the HIT motif, HIT1 subdomain, HIT2 subdomain, auxiliary domain, and disordered regions are colored magenta, green, cyan, pink, and gray, respectively. (B) Apa2 complexed with Ap_4_A (yellow). Protein residues are color-coded as in (A). (C) *E. coli* GalT complexed with UDP-Gal (yellow). Residues in the HIT motif, other chain A residues, and residues from the neighboring subunit (chain B, indicated by a prime mark) are shown in magenta, blue, and gray, respectively. Hydrogen bonds are indicated by yellow dotted lines.

Recognition of the hydroxy groups of the GalNAc moiety was mediated by Q290 in the HIT motif, as well as by residues in the auxiliary domain (K412, E414, E420, and P415; Fig. 6A, right). Although I419 did not directly interact with the substrate, it was located close to the C6 position of GalNAc (with a carbon-carbon distance of 3.9 Å). The *N*-acetyl group was accommodated in a hydrophobic pocket formed by F176, M179, and A182. These residues are located in the helix predicted to cover the active site and in the long disordered region (Fig. 5). Thus, highly flexible regions are also likely to contribute to the preference of BlGalT2 for GalNAc and GlcNAc-containing substrates.

We then compared the active site of BlGalT2 with those of the well-characterized Apa2 and EcGalT. In Apa2, the substrate (Ap_4_A) is recognized by residues from HIT1 (N92), HIT2 (N148), and the auxiliary domain (N277, Fig. 6B). In EcGalT, residues from the neighboring subunit (F312’, V314’, Y316’, and E137’) mainly recognize the Glc moiety of the substrate, UDP-Glc (Fig. 6C). Therefore, BlGalT2, which recognizes the sugar moiety of the substrate via its auxiliary domain, possesses a substrate recognition mechanism distinct from that of class I GalTs. Although residues involved in substrate recognition are poorly conserved among Apa2, EcGalT, and BlGalT2, the serine residue involved in the recognition of the α-phosphate group (donor phosphate) targeted by nucleophilic attack (S156 in Apa2 and S161 in EcGalT) is conserved in BlGalT2 (S282). In addition, both the hydrogen bond between the two histidines in the HIT motif (H159-H161 in Apa2 and H164-H166 in EcGalT) and the position of the third glutamine residue (Q163 in Apa2 and Q168 in EcGalT) are conserved. The third glutamine forms a hydrogen bond with the β-phosphate group (the acceptor phosphate) in Apa2 and with the α-phosphate group in EcGalT, and has been suggested to stabilize the reaction intermediate [32,34]. Although metal binding to H159 has not been confirmed in Apa2 [32], Zn²⁺ is coordinated by the first HIT motif histidine in all reported crystal structures of class I GalTs [28,29,34–36], including At5g18200 [33] and EcGalT (Fig. 4C,D and Fig. 6C). In the reaction mechanisms proposed for Apa2 and EcGalT, however, the metal does not directly participate in catalysis [32,35]. In EcGalT, Zn^2+^ has instead been proposed to stabilize the second, nucleophilic histidine [27]. These observations suggest that BlGalT2 follows a reaction mechanism similar to that proposed for Apa2 and EcGalT.

### Mutational analysis

To examine the contribution of individual amino acid residues in the binding sites of UDP-GlcNAc and Zn^2+^ inferred from the crystal and predicted structures, we generated alanine-substitution mutants by site-directed mutagenesis and measured their activities (Table 4). No activity was detected for the H286A, H288A, and Q290A mutants in the HIT motif, whereas the D297A mutant, located between the two histidines, retained approximately half of the wild-type activity. No activity was detected for either of the two cysteine mutants (C195A and C198A), which were predicted to coordinate Zn^2+^ together with H286. All mutants of the residues predicted to form hydrogen bonds with UDP-GlcNAc (Y233A, K412A, and E420A) showed no detectable activity. In addition, the mutant of a conserved asparagine residue (N273A) lost activity, indicating that N273 is also involved in substrate binding. However, the I419A, whose side chain forms a hydrophobic environment near the GlcNAc moiety, retained approximately 1.2% of wild-type activity. These results support the validity of the predicted substrate-bound structure of BlGalT2.

**Table 4.** Activity of site-directed mutants.

| Enzyme | Specific activity (U/mg) |
| --- | --- |
| Wild type | 0.61 ± 0.10 |
| C195A | N.D. |
| C198A | N.D. |
| Y233A | N.D. |
| N273A | N.D. |
| H286A | N.T. |
| D287A | 0.30 ± 0.02 |
| H288A | N.D. |
| Q290A | N.D. |
| K412A | N.D. |
| I419A | 0.074 ± 0.03 |
| E420A | N.D. |

N.D., not detected (< 0.005 U/mg). N.T., not tested due to protein aggregation during the enzyme assay. Protein concentrations of the enzyme stock solution used for the assay was 0.10 mg/mL for the wild-type, D278A and I419A, and 10 mg/mL for other mutants.

## DISCUSSION

### Reaction mechanism

Based on the findings of this study, we propose a reaction mechanism for the uridylyl group transfer from UDP-Glc(NAc) to Gal(NAc)-1P catalyzed by BlGalT2 (Fig. 7). The side chain of H288, which nucleophilically attacks the α-phosphate of the nucleotide sugar substrate, is stabilized by a hydrogen bond with the main chain carbonyl of H286. Coordination of Zn^2+^ by the side chain of H286 also contributes indirectly to this nucleophile stabilization. Formation of the covalent intermediate (E–UMP) is accompanied by cleavage of the phosphodiester bond in the nucleotide sugar substrate. This reaction is facilitated by the hydrogen bond and salt bridge formed by the side chains of Q290 and K173 with the β-phosphate, as well as an oxyanion hole for the α-phosphate formed by the main chain amide nitrogens of S282 and V283. After the phosphorylated sugar product dissociates from the active site and the acceptor substrate enters, the subsequent reaction proceeds. Nucleophilic attack by the acceptor on the phosphate group of E–UMP leads to cleavage of the covalent bond in the nucleotidylated histidine, and the formation of a new phosphodiester bond produces the nucleotide sugar product, thereby completing the transfer reaction.

**Figure 7.**
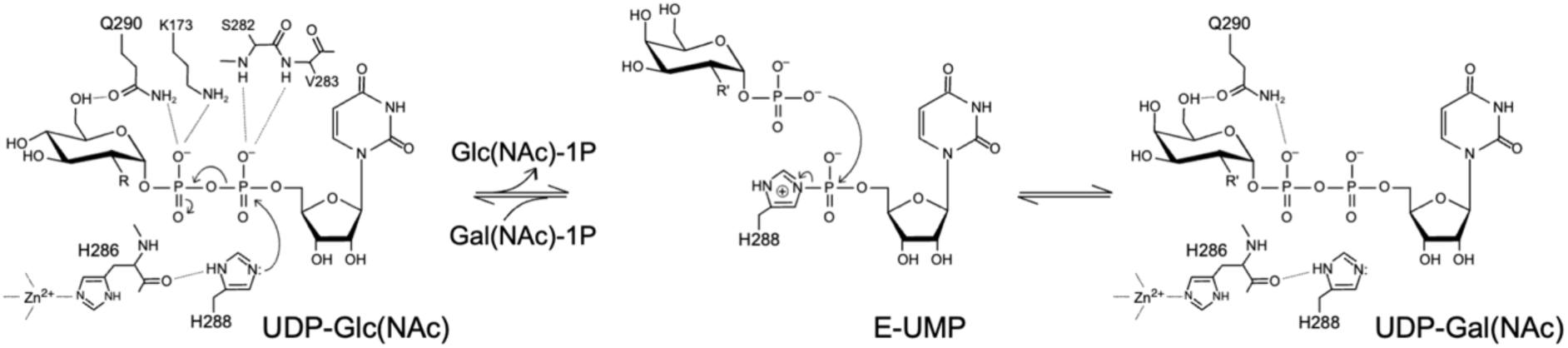
Proposed reaction mechanism of BlGalT2. R = −OH for UDP-Glc and Glc-1P; R = - NHCOCH_3_ for UDP-GlcNAc and GlcNAc-1P.

### Active site architecture and substrate preference of BlGalT2

Our kinetic and structural analyses revealed that BlGalT2 exhibits a significantly higher affinity for GalNAc-1P than for Gal-1P (Table 2), and its active site features a hydrophobic pocket suitable for the *N*-acetyl group of the substrate (Fig. 6A, right). Among the class I and class II GalTs in *B. bifidum*, BbGalT1 exhibits activity only toward Gal-1P (apparent *K*_m_ = 0.29 mM), whereas BbGalT2 shows a lower apparent *K*_m_ for GalNAc-1P (0.065 mM) than for Gal-1P (0.61 mM) [16]. De Bruyn et al. [16] suggested that BbGalT1 is responsible for the metabolism of Gal-1P, whereas BbGalT2 is involved in the GNB/LNB pathway of HMO degradation. Our study provides the structural basis for the selective utilization of the two GalT types across metabolic pathways, governed by the unique substrate specificity of class II enzymes.

### A class II GalT scaffold distinct from class I GalTs

The structural elucidation of BlGalT2 offers the first three-dimensional insights into class II GalT, revealing an architectural topology distinct from that of class I GalTs, despite their identical UDP-sugar transferase activity. Although BlGalT2 retains the characteristic HIT-family catalytic core, its closest structural relatives are monomeric Ap_4_A phosphorylases (such as Apa2) rather than class I GalTs, illustrating that activity-based classification obscures the structural divergence within the GalT enzyme family. Most notably, its solution behavior and crystal structure confirm that BlGalT2 functions as a monomer, in contrast to canonical class I GalTs that strictly require dimerization for a neighboring subunit to complete the substrate-binding pocket. BlGalT2 circumvents this requirement through its extended auxiliary domain, which supplies key sugar-recognizing residues (K412, I419, E420, and I426) to form a complete, self-contained active site. This structural feature represents a defining adaptation of class II GalTs. Additionally, a disordered linker adjacent to the proposed nucleotide-binding region may play a functional role, although its exact conformational dynamics remain unresolved in the present crystal structure.

### Structural features and possible molecular evolution of the enzymes in the GNB/LNB pathway

The GNB/LNB pathway metabolizes host-derived LNB and GNB through the following five steps that are essentially conserved in infant-associated bifidobacteria: cellular import by a specific transporter (GL-BP), phosphorolysis (GLNBP), C1-phosphorylation of *N*-acetylhexosamines (NahK), C4-epimerization of nucleotide sugars (bGalE), and uridylyl transfer between nucleotide sugars and phosphorylated sugars (BlGalT2) [8]. Because structural information has previously been reported for the first four steps [6,10–12], the determination of the BlGalT2 structure therefore completes the structural framework of this pathway. The three downstream enzymes (NahK, bGalE, and BlGalT2) are functionally related to the enzymes in the conventional galactose-metabolizing Leloir pathway, namely galactokinase (GalK), UDP-Gal 4-epimerase (GalE), and class I Gal-1P uridylyltransferase (GalT) [37,38]. A key functional difference distinguishing the three GNB/LNB pathway enzymes from those in the Leloir pathway is their ability to act on *N*-acetylated sugars. Indeed, NahK, bGalE, and BlGalT2 can be used to efficiently produce GalNAc from GlcNAc in a one-pot reaction [39]. Among them, NahK is structurally distinct from GalK but closer to homoserine kinases and aminoglycoside phosphotransferases, probably due to the requirement for recognizing the sugar *N*-acetyl moiety during the large open-to-closed domain movement accompanying catalysis [11]. In contrast, the structure of bGalE closely resembles that of canonical GalEs because a substitution at a specific amino acid position can extend its substrate preference from UDP-Glc/Gal to UDP-GlcNAc/GalNAc [12]. In this study, BlGalT2 provides a particularly clear example of specialized molecular evolution for the utilization of *N*-acetylated host glycans based on a unique class II GalT scaffold distinct from conventional class I GalTs. Thus, while the GNB/LNB pathway represents a variation of the Leloir pathway, its enzymes have evolved unique structural features that tailor them to the glycan environment of the infant gut microbiome. Obtaining experimentally determined ligand-bound three-dimensional structures of BlGalT2 or other class II GalTs in the future will further elucidate the atomic-level mechanism of *N*-acetylhexosamine recognition.

## ACKNOWLEDGMENTS

We thank Dr. Takatoshi Arakawa for his technical help and insightful discussions; Profs. Takane Katayama, Hisashi Ashida, and Kenji Yamamoto for insightful discussions; Dr. Toma Kashima for assistance on protein structure determination; the staff of KEK-PF and SPring-8 for help with X-ray data collection; and the Organization for Open Facility Initiatives at University of Tsukuba for ICP-MS measurements. This work was supported by the Program for the Promotion of Basic Research Activities for Innovative Bioscience, Japan (PROBRAIN to M.K., M.N., and S.F.), Science and Technology Research Promotion Program for Agriculture, Forestry, Fisheries and Food Industry (25010A to M.K., M.N., and S.F.), and Grant-in-Aid for JSPS Fellows (to M.S.), and in part by JSPS-KAKENHI (24380053, 26660083, 15H02443, 23H00322, and 24H02269 to S.F.) and Platform Project for Supporting Drug Discovery and Life Science Research (Basis for Supporting Innovative Drug Discovery and Life Science Research (BINDS)) from AMED under Grant Number JP21am0101071.

## Abbreviations

ABC: ATP-binding cassette
BbGalT1: *B. bifidum* class I GalT
BbGalT2: *B. bifidum* class II GalT
bGalE: *B. longum* UDP-glucose 4-epimerase
BlGalT2: *B. longum* class II GalT
EcGalT: *E. coli* GalT
EDTA: ethylenediaminetetraacetic acid
E–UMP: enzyme-uridine monophosphate intermediate
GalT: UDP-glucose—hexose-1-phosphate uridylyltransferase
GLNBP: GNB/LNB phosphorylase
GL-BP: solute-binding protein of GNB/LNB-specific ABC-type transporter
GNB: galacto-*N*-biose
hGalT: human GalT
HIT: histidine triad
HMOs: human milk oligosaccharides
ICP-MS: inductively coupled plasma mass spectrometry
LNB: lacto-*N*-biose I
NahK: *N*-acetylhexosamine 1-kinase
PaGalT: *P. aerophilum* GalT
RMSD: root-mean-square deviations
SeMet: selenomethionine.

## Supplemental Data

**Table S1.**
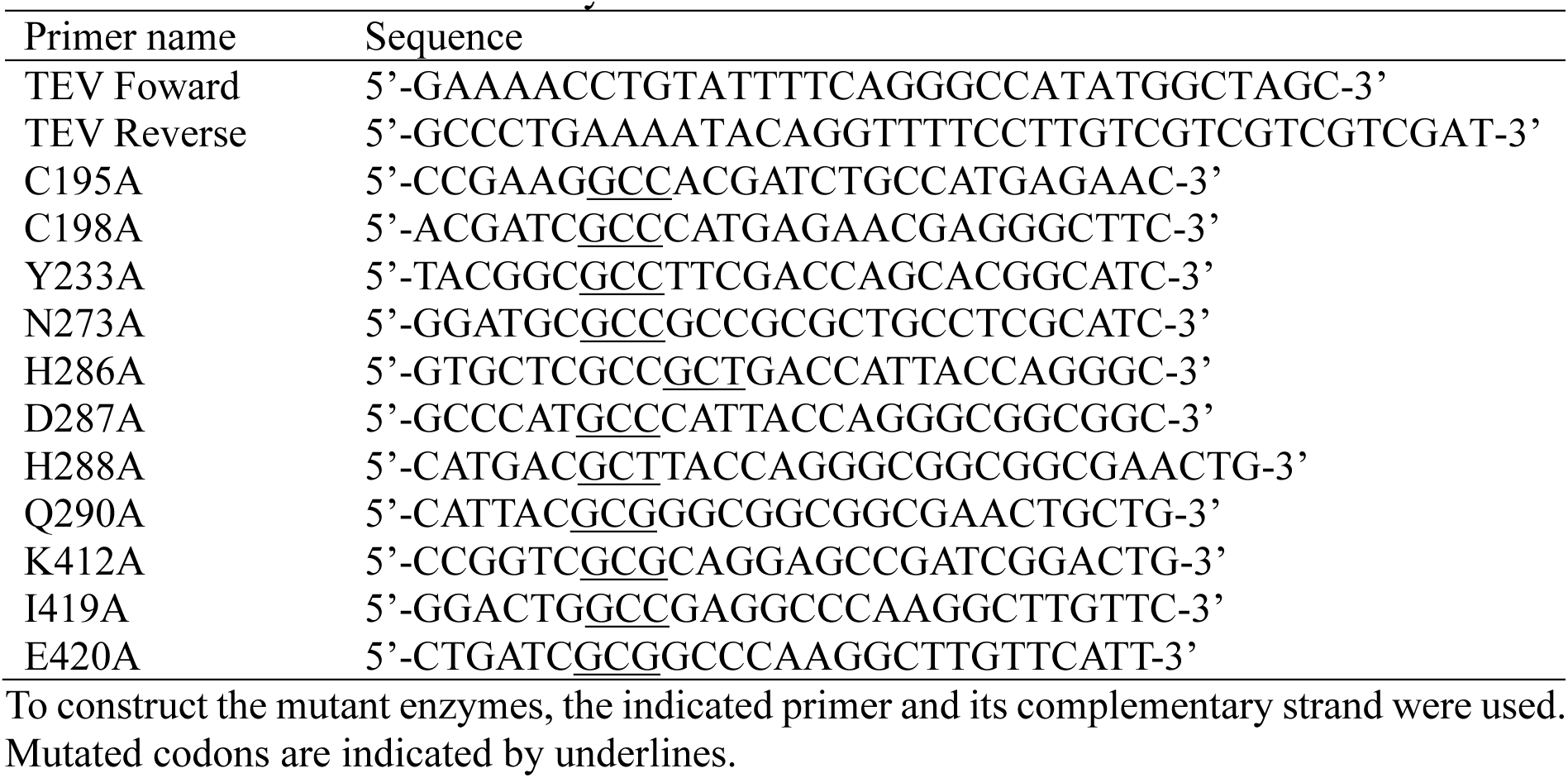
Primers used in this study.

**Table S2.** Metal quantification data of purified BlGalT2.

| **Metal ion** | **Protein solution** | **Buffer solution** | **Protein–Buffer** | **Metal content (mol/mol)** |
| --- | --- | --- | --- | --- |
| **Dilution ×1 (0.075 mg)** |  |  |  |  |
| Mg<sup>2+</sup> | 3.97 | 4.16 | -0.19 | (-0.03) |
| Ca<sup>2+</sup> | 64.89 | 91.42 | -26.53 | (-2.26) |
| Mn<sup>2+</sup> | 1.24 | 0.26 | 0.98 | 0.07 |
| Fe<sup>2+</sup> | 24.56 | 20.84 | 3.71 | 0.25 |
| Co<sup>2+</sup> | 0.01 | 0.01 | 0.00 | 0.00 |
| Ni<sup>2+</sup> | 1.53 | 0.82 | 0.71 | 0.04 |
| Cu<sup>2+</sup> | 2.01 | 1.94 | 0.07 | 0.00 |
| V<sup>2+</sup> | 0.02 | 0.02 | 0.01 | 0.00 |
| Cr<sup>2+</sup> | 0.22 | 0.21 | 0.01 | 0.00 |
| Zn<sup>2+</sup> | 19.93 | 3.38 | 16.55 | 1.19 |
| **Dilution ×1/5 (0.015 mg)** |  |  |  |  |
| Mg<sup>2+</sup> | 1.07 | 1.22 | -0.14 | (-0.11) |
| Ca<sup>2+</sup> | 15.66 | 22.72 | -7.05 | (-3.00) |
| Mn<sup>2+</sup> | 0.25 | 0.06 | 0.19 | 0.07 |
| Fe<sup>2+</sup> | 5.04 | 4.36 | 0.69 | 0.23 |
| Co<sup>2+</sup> | 0.03 | 0.00 | 0.03 | 0.01 |
| Ni<sup>2+</sup> | 0.31 | 0.17 | 0.14 | 0.04 |
| Cu<sup>2+</sup> | 0.72 | 0.69 | 0.03 | 0.01 |
| V<sup>2+</sup> | 0.00 | 0.00 | 0.00 | 0.00 |
| Cr<sup>2+</sup> | 0.05 | 0.06 | -0.02 | -0.01 |
| Zn<sup>2+</sup> | 4.09 | 0.89 | 3.20 | 0.94 |
| **Dilution ×1/25 (0.003 mg)** |  |  |  |  |
| Mg<sup>2+</sup> | 0.41 | 0.28 | 0.13 | 0.50 |
| Ca<sup>2+</sup> | 7.20 | 6.79 | 0.41 | 0.87 |
| Mn<sup>2+</sup> | 0.06 | 0.01 | 0.05 | 0.09 |
| Fe<sup>2+</sup> | 1.24 | 0.98 | 0.26 | 0.43 |
| Co<sup>2+</sup> | 0.00 | 0.00 | 0.00 | 0.00 |
| Ni<sup>2+</sup> | 0.07 | 0.04 | 0.03 | 0.04 |
| Cu<sup>2+</sup> | 0.16 | 0.14 | 0.02 | 0.03 |
| V<sup>2+</sup> | -0.01 | 0.01 | -0.02 | -0.03 |
| Cr<sup>2+</sup> | 0.01 | 0.01 | 0.00 | 0.00 |
| Zn<sup>2+</sup> | 0.87 | 0.16 | 0.71 | 1.04 |

The dilution fold and protein amount of the three measurements are shown. Values in the columns of protein solution, buffer solution, and protein-buffer are raw intensity values of ICP-MS measurements. The metal content was calculated from the protein content and mass and the atomic mass of each metal.

**Table S3.** Crystallographic data statistics of BlGalT2.

| Data set | Se-Met labeled | Native |
| --- | --- | --- |
| **Data collection** |  |  |
| Beamline | KEK-PF BL17A | SPring-8 BL26B1 |
| Wavelength (Å) | 0.97880 | 1.00000 |
| Space group | $P2_12_12_1$ | $P2_12_12_1$ |
| Unit cell (Å) | $a = 84.29, b = 92.53, c = 159.45$ | $a = 84.79, b = 94.42, c = 160.51$ |
| Resolution (Å) | 49.09–2.70 (2.83–2.70) | 47.21–2.25 (2.31–2.25) |
| Total reflections | 931202 (124515) | 453991 (32047) |
| Unique reflections | 35033 (4572) | 61778 (4421) |
| $R_{\text{merge}}$ | 0.240 (1.452) | 0.133 (1.868) |
| $R_{\text{meas}}$ | 0.244 (1.479) | 0.144 (2.012) |
| $R_{\text{pim}}$ | 0.047 (0.282) | 0.053 (0.742) |
| Mean $I/\sigma(I)$ | 15.8 (3.1) | 8.3 (1.2) |
| $CC_{1/2}$ | 0.998 (0.927) | 0.997 (0.819) |
| Completeness (%) | 100.0 (100.0) | 99.8 (97.6) |
| Multiplicity | 26.6 (27.2) | 7.3 (7.2) |
| Anomalous completeness (%) | 100.0 (100.0) |  |
| Anomalous multiplicity | 13.9 (14.0) |  |
| **Refinement** |  |  |
| Resolution (Å) |  | 47.25–2.25 |
| No. of reflections |  | 61648 |
| $R_{\text{work}}/R_{\text{free}}$ | | 0.236/0.276 |
| Number of atoms |  |  |
| Amino acids |  | 7013 |
| Ions |  | 2 |
| Waters |  | 192 |
| B-factors (Å<sup>2</sup>) |  |  |
| Amino acids<sup>a</sup> |  | 38.9/41.4 |
| Ions |  | 40.0 |
| Waters |  | 31.6 |
| MolProbity clashscore |  | 3.98 |
| MolProbity score |  | 1.85 |
| RMSD from ideal values |  |  |
| Bond lengths (Å) |  | 0.0069 |
| Bond angles (°) |  | 1.578 |
| Ramachandran plot (%) |  |  |
| Favored |  | 96.35 |
| Allowed |  | 3.08 |
| Outlier |  | 0.57 |
| PDB code |  | 45PI |

Values in parentheses are for the highest resolution shell.
<sup>a</sup> Atoms in chain A/chain B are shown.

**Figure S1.**
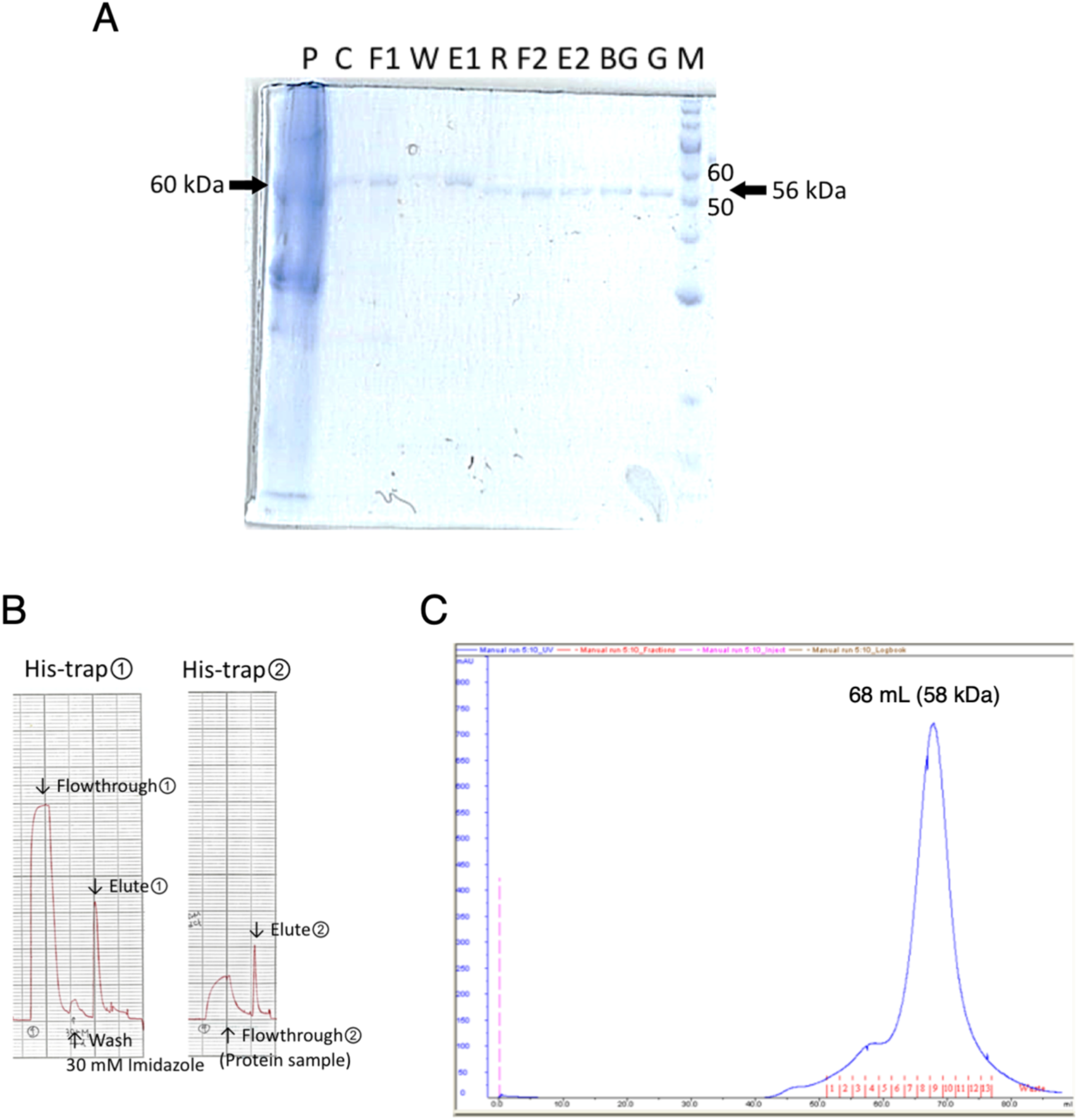
Purification of BlGalT2. (A) SDS-PAGE of fractions from each purification step. Symbols: P and C, pellet and crude supernatant after cell sonication and centrifugation; F1, W, and E1, flow-through, wash, and eluate fractions from the first HisTrap column purification; R, reaction mixture after TEV protease treatment; F2 and E2, flow-through and eluate fractions of the second HisTrap column purification; BG and G, samples before and after gel filtration purification; M, molecular weight marker. (B) Chromatograms of the first (left) and second (right) HisTrap column purification steps. (C) Chromatogram of gel filtration purification using a HiLoad 16/600 Superdex 200 pg column.

**Figure S2.**
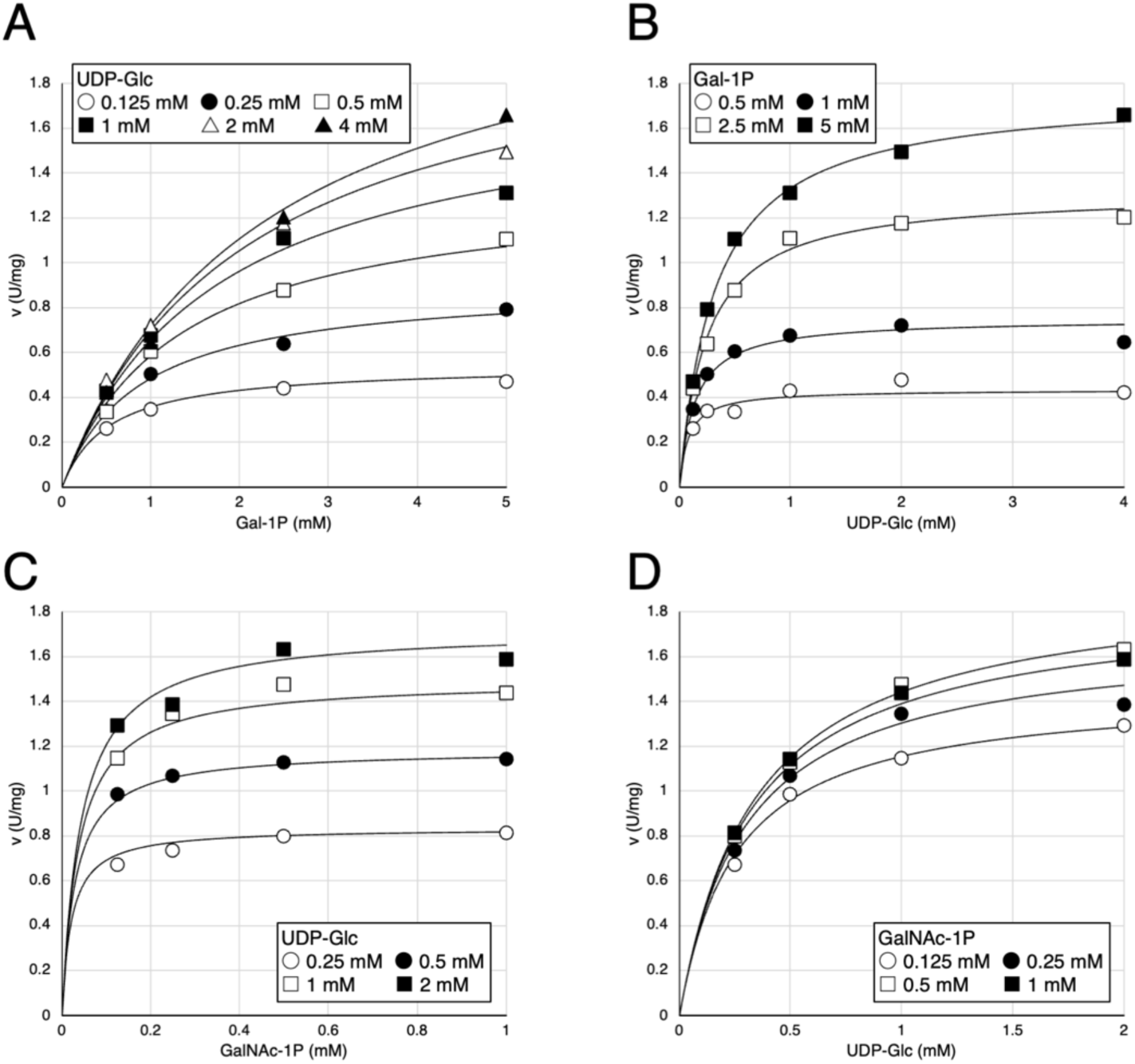
Substrate-velocity (*v* vs [*S*]) plots of BlGalT2 for the reactions with UDP-Glc and Gal-1P (A and B), and UDP-Glc and GalNAc-1P (C and D). The data were fitted to the Ping-Pong Bi-Bi kinetic equation: *v* = *V*_max_[A][B]/(*K*_mB_[A] + *K*_mA_[B] + [A][B]). The kinetic parameters are shown in Table 2.

**Figure S3.**
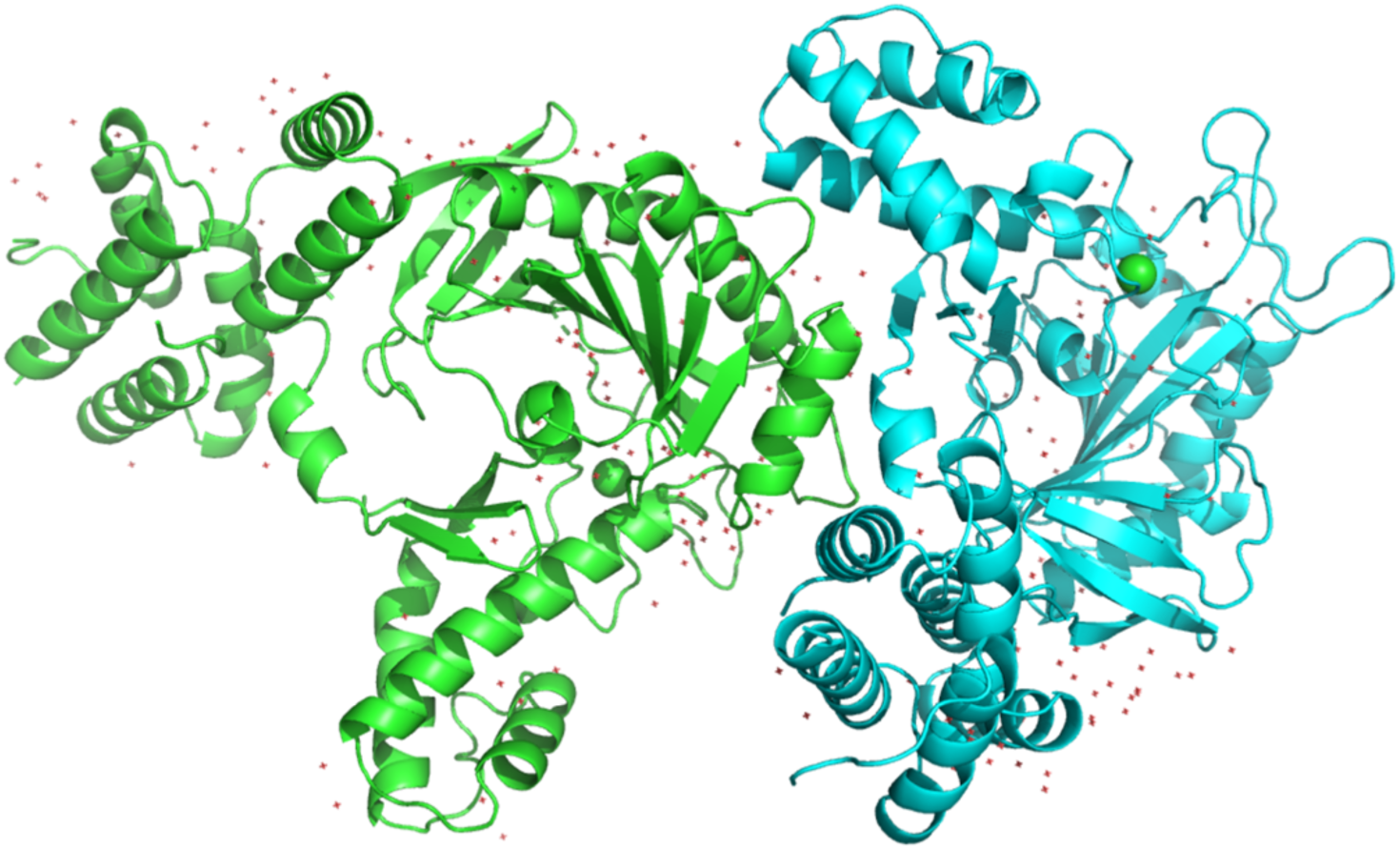
Asymmetric unit of BlGalT2 crystal. Chains A and B are shown in green and cyan, respectively. Water molecules and Cl atoms are shown as red crosshairs and green spheres, respectively.

**Figure S4.**
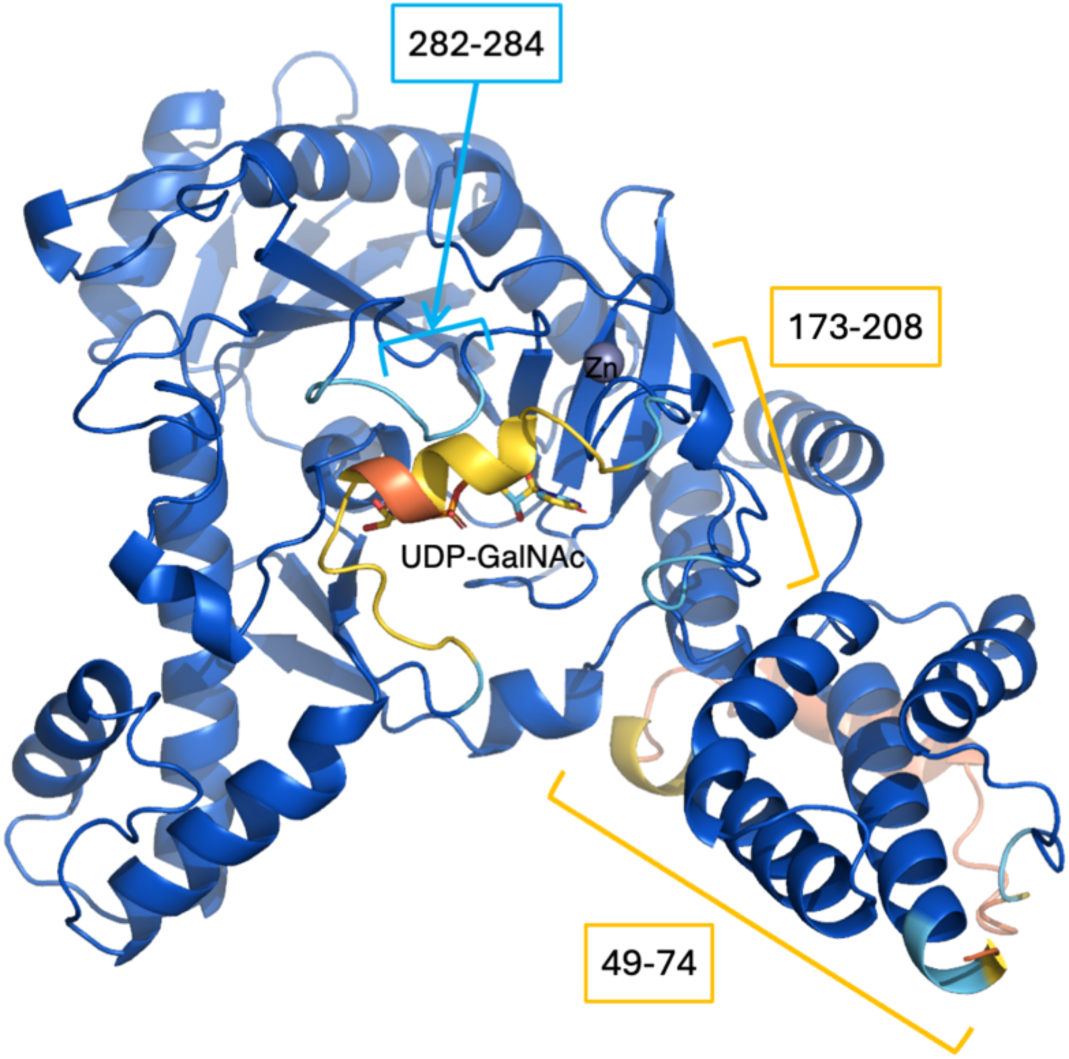
Alphafold-predicted structure of BlGalT2 colored by predicted local distance difference test (pLDDT) score: dark blue (>90), light blue (>70), yellow (>50), and orange (≤ 50). The three disordered regions in the crystal structure are indicated.

